# The contributions of extracellular matrix and sarcomere properties to passive muscle stiffness in cerebral palsy

**DOI:** 10.1101/2021.06.17.447742

**Authors:** Ryan N. Konno, Nilima Nigam, James M. Wakeling, Stephanie A. Ross

**Affiliations:** Department of Mathematics, Simon Fraser University; Department of Biomedical Physiology and Kinesiology, Simon Fraser University

## Abstract

Cerebral palsy results from an upper motor neuron lesion and has significant effects on skeletal muscle stiffness throughout the body. The increased stiffness that occurs is partly a result of changes in the microstructural components of muscle. In particular, alterations in extracellular matrix, sarcomere length, fibre diameter, and fat content have been reported; however, experimental studies have shown wide variability in the degree to which each component is altered. Many studies have reported alterations in the extracellular matrix, while others have reported no changes. A consistent finding throughout the literature is increased sarcomere length in cerebral palsy muscle. Often more than one component is altered, making it difficult to determine the individual effects on stiffness. The purpose of this study is to use a modeling approach to isolate individual effects of microstructural alterations that typically occur during cerebral palsy on whole muscle behavior; in particular, the extracellular matrix volume fraction, stiffness, and sarcomere length. These microstructural effects can be captured using a three dimensional model of muscle. We found that the extracellular matrix volume fraction has a larger effect on stiffness compared to sarcomere length, even when coupled with decreased extracellular matrix stiffness. Additionally, the effects of sarcomere length in passive stiffness are mitigated by the increased extracellular matrix volume fraction. Using this model, we can achieve a better understanding of the possible combinations of microstructural changes that can occur during cerebral palsy. Developing these insights into diseased muscle tissue will help to direct future clinical and experimental procedures.

## 1 Introduction

Cerebral palsy (CP) results from an upper motor neuron lesion and has a significant effect on the musculoskeletal system. It develops during early childhood and leads to muscle alterations including contracture, which is the chronic shortening of muscle. Contracture results in muscle that cannot be stretched throughout its typical range of motion due to an increase in stiffness, and this has substantial effects on the ability of muscle to generate force and reduces daily functioning. CP will affect individuals differently, and the changes that can occur will vary depending on the location of the muscle [de Bruin et al., 2014, Lieber and Fridén, 2019]. This variability increases the difficulty in quantifying the amount and types of changes that occur during CP. However, despite the variability, alterations in the microstructural properties of skeletal muscle are commonly observed [Tisha et al., 2019], which will have a significant effect on whole muscle behavior, including force production and movement.

There are many structural differences comparing CP muscle to typically developed (TD) muscle, including changes in fat content [D’Souza et al., 2020, Ohata et al., 2009], extracellular matrix (ECM) stiffness [Lieber et al., 2003, Smith et al., 2011], amount of ECM [Lieber et al., 2003, Smith et al., 2011], fascicle length [Mohagheghi et al., 2008], fibre diameter [Mathewson et al., 2014a], and sarcomere length [Lieber and Fridén, 2002, Mathewson et al., 2014a, Smith et al., 2011]. Experimental studies have investigated CP muscle stiffness *in vivo* and have found stiffer tissue compared to TD muscle using shear wave elastography [Brandenburg et al., 2016, Lee et al., 2016] and through measuring joint movement [Barber et al., 2011, van der Krogt et al., 2016]. However, these methods are unable to capture the underlying causes of this increased stiffness.

The exact microstructural changes that alter whole muscle stiffness have yet to be fully understood, as the extent of measured changes varies between studies [Lieber and Fridén, 2019]. For example, Smith et al. [2011] performed passive mechanical experiments on both muscle fibre bundles and single fibres extracted from CP and TD muscle. They found that CP muscle had longer *in vivo* sarcomere lengths and increased fibre bundle, but not fibre, stiffness, which suggests that the changes in muscle stiffness are due to alterations in the ECM. Another study by Mathewson et al. [2014a], which used a similar experimental protocol as Smith et al. [2011], also showed increased *in vivo* sarcomere lengths. However, the authors demonstrated a difference in the single fibre stiffness and not the fibre bundles, suggesting that there is not a significant effect from the ECM, and that any alterations to passive stiffness occur on the muscle fibre level. Other studies have reported that CP muscle has a greater accumulation of fibrotic tissue, and potentially even results in a ECM with a larger volume fraction but compromised stiffness [Lieber et al., 2003]. Additionally, Booth et al. [2001] suggested that collagen plays a role in the increased muscle stiffness that is observed in CP. However, it has been observed that fibrosis does not always alter the stiffness of muscle [Smith and Barton, 2014], and so there may be an effect from the stiffness and structure of the ECM. Sarcomere length is a commonly observed alteration in CP muscle, and has been said to have a large effect on active muscle mechanics [Lieber and Fridén, 2002, Mathewson et al., 2014b, Smith et al., 2011]. While many changes have been observed in CP muscle, the changes that are most common between studies are changes in ECM volume fraction, ECM stiffness, and sarcomere length. However, the individual roles of the ECM properties and sarcomere length in passive muscle stiffness have yet to be fully understood.

The purpose of this study was to determine which microstructural change occurring during CP has the largest contribution to whole muscle stiffness. In particular, whether the ECM, through changes in volume fraction or stiffness, or the sarcomere, through increases in length that result in an increased passive response from the titin, will have the greatest influence on passive whole muscle behavior. It is difficult to investigate this using experimental techniques due to the variability in muscle composition that occurs between individuals. However, using a modeling approach, we investigated the influence of the microstructural components on passive muscle stiffness. We utilized a three dimensional continuum model of skeletal muscle, developed in previous studies [Konno et al., 2021, Rahemi et al., 2014, Ross et al., 2018, Wakeling et al., 2020], which can be modified to incorporate the effects of ECM and passive fibre properties on whole muscle mechanics. By modifying the material properties in the muscle, we investigated changes that occur during CP to understand how each component contributes to muscle stiffness.

## 2 Methods

### 2.1 Computational Model

In this study, we utilized a continuum mechanical model of muscle as a fibre-reinforced composite biomaterial. This model is split into a three dimensional isotropic base material with one dimensional fibres running along the length of the muscle, making the composite material anisotropic. We characterized the passive mechanical behavior of muscle in terms of its stress-strain behavior, which is the relationship between the stress applied to the muscle and the strain experienced. A precise formulation of the model is described in Wakeling et al. [2020] and Konno et al. [2021]. We used a finite element method to solve the continuum model that was implemented using an open source finite element library deal.II [Arndt et al., 2017].

The total stress response from the muscle (*σ*_muscle_) was broken down into contributions from the base material (*σ*_base_) and fibre (*σ*_fibre_) components. Here the base material encompasses effects from extracellular matrix and cellular components, including satellite cells, neuron bodies, and capillaries, while the fibre component runs along the length of the muscle and contains the passive effects from titin and active effects from the contractile elements. To investigate the role of CP on whole muscle stiffness, considering the individual effects from the ECM and cellular components is necessary. To do this, we let *α* be the volume fraction of the ECM, which can increase during muscle fibrosis [Lieber et al., 2003], and so 1 − *α* is the volume fraction of the cellular component. We also introduced a parameter *s*_ECM_, which is a stiffness factor multiplying the stiffness of ECM. A larger *s*_ECM_ corresponds to a stiffer ECM, which can occur as a result of structural changes while the ECM volume fraction remains the same [Gillies et al., 2016]. Meanwhile, a smaller value of *s*_ECM_ results in an ECM with decreased stiffness, which can occur as a result of an ECM with compromised structure [Lieber et al., 2003]. Therefore, the total stress response from the base material is given by

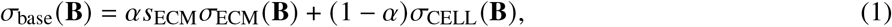

where **B** is a strain tensor measuring the deformation in the material. *σ*_ECM_ (**B**) corresponds to the stress response from the ECM, while *σ*_CELL_ (**B**) is the stress response from the cellular component. To ensure the volume of the muscle remains nearly constant, the bulk modulus for the cellular component is chosen to be 1 × 10^7^ Pa, while the bulk modulus of the ECM component was set to 1 × 10^6^ Pa [Konno et al., 2021]. The exact form of the stress-strain response is given in Konno et al. [2021].

The other alteration typically observed in CP is an increase in *in vivo* sarcomere length [Mathewson et al., 2014b, Smith et al., 2011], which alters the passive muscle stiffness by stretching the titin protein. The *in vivo* length of the sarcomeres can be defined as the length of the sarcomeres when the whole muscle is at its resting length. This change in length also decreases the contractile force produced when the muscle is active by reducing the number of attached actin-myosin crossbridges (Figure 1). We modelled this using a dimensionless parameter, *c*_sarco_, which corresponds to a shift in the passive force length curve of the sarcomeres

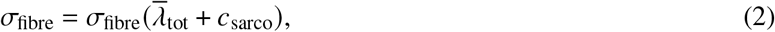

where 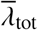 is the total average stretch of the fibres over the muscle volume. It is important to note that, while the fibre component of the muscle depends on *c*_sarco_, the intrinsic stress response from the base material, *σ*_base_ (**B**), only depends on the deformation and stretch of the muscle, and not *c*_sarco_. At a value of *c*_sarco_ = 0.0, the behavior of the sarcomere is the same as that of TD muscle. Increasing values of *c*_sarco_ results in longer lengths of the sarcomeres given by

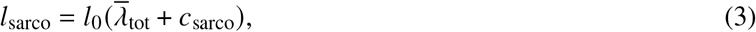

where *l*_sarco_ is the new length of the sarcomere and we assume *l*_0_ = 2.2*μ*m is the optimal length of a sarcomere [Burkholder and Lieber, 2001]. This will vary depending on the value of *c*_sarco_. The fibres in the model are based off the one dimensional Hill type model [Hill, 1938, Zajac, 1989] and are described in Wakeling et al. [2020].

**Figure 1:**
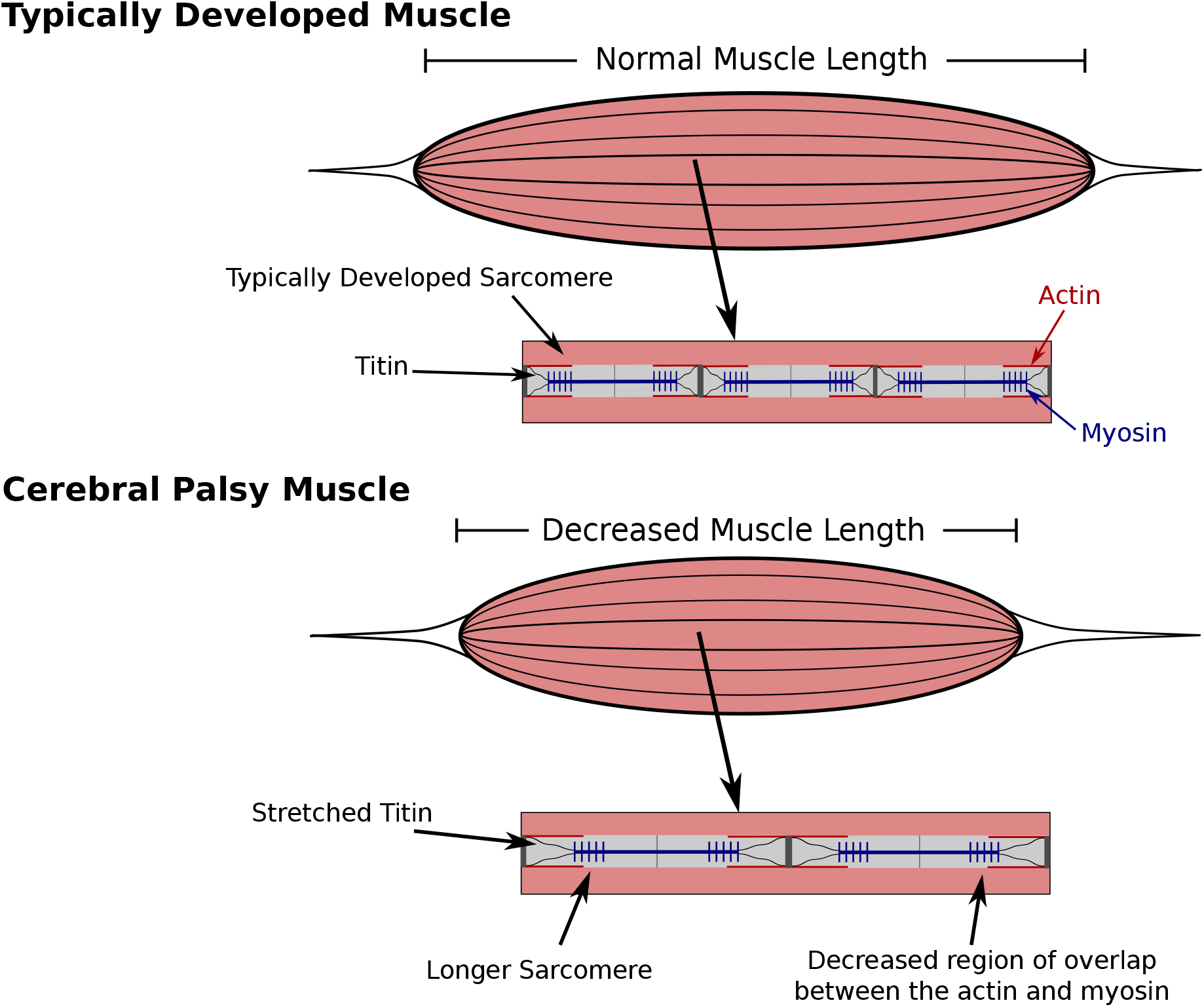
Comparison of typically developed (TD) and cerebral palsy (CP) muscle. CP results in contracture, which is the chronic shortening of muscle, decreasing muscle length relative to TD muscle (not investigated in this study). Longer sarcomere lengths relative to the rest of the muscle have also been observed compared to TD muscle [Mathewson et al., 2014b, Smith et al., 2011]. There is an increase in passive forces due to the increased stretch in the titin proteins. The longer sarcomeres lengths in muscle with CP reduces the regions of overlap of the actin and myosin filaments, which results in decreased contractile forces.

### 2.2 Whole Muscle Experiments

To investigate the passive effects of *α*, *s*_ECM_, and *c*_sarco_ on skeletal muscle, we constructed a rectangular block of muscle with dimensions 3cm × 1cm × 1cm. These dimensions, while not the same dimensions as muscle affected by CP, sufficiently capture the behavior of muscle on a macroscopic scale. Using a block geometry reduces the need to consider the additional effects from geometry, aponeurosis, and pennation angle that has been shown to affect muscle behavior [Wakeling et al., 2020], and instead allowed us to isolate the effects due to CP. This geometry had the additional benefit of computational simplicity, and these geometries have been shown to exhibit similar qualitative behavior to a block of muscle in real muscle tissue [Wakeling et al., 2020]. To compare the passive behavior of the model to experimental data, we performed stress-strain tests. This involved constraining one end face of the model from movement in all directions, while a normal stress was applied to the opposite face stretching the muscle. In addition to the stress-strain tests, we investigated the stiffness of the muscle to compare with experimental studies (eg. Mathewson et al. [2014b], Smith et al. [2011]). *α*, *s*_ECM_, and *c*_sarco_ each have an individual contribution to the overall stiffness of muscle, and to investigate the stiffness in the model, the modulus of the muscle material was calculated during the stress-strain experiments. The modulus of the muscle (in Pa) was calculated as the slope of the tangent line to the overall stress-stretch relationship and calculated at individual stretches. This was done by performing a nonlinear least-squares fit of a cubic polynomial to the overall stress-strain data in the longitudinal direction.

TD muscle has been observed to have a value for *α* between 0.02-0.10 [Binder-Markey et al., 2020], while larger volume fractions (*α* ≈ 0.6) have been observed for fibrotic tissue [Lieber et al., 2003, Smith and Barton, 2014]. Experimental studies have only found an increase in *in vivo* sarcomere lengths [Lieber and Fridén, 2002, Mathewson et al., 2014a, Smith et al., 2011], so we varied *c*_sarco_ from 0 to 0.75. This corresponded to a longer length of the sarcomeres relative to the muscle fibres, which has been observed in the literature [Lieber and Fridén, 2002, Mathewson et al., 2014b, Smith et al., 2011]. Experiments are often performed on advanced cases of CP, so larger sarcomere lengths have been reported (*c*_sarco_ > 0.75); however, in this study, we considered smaller sarcomere lengths to represent less severe cases of CP. It is likely that muscle altered by CP does not always have such a substantial increase in sarcomere length. The final parameter that was manipulated in the model is *s*_ECM_. While the stiffness of muscle can vary depending on the type of muscle, the properties of the base material, including the effects from the ECM, represent and have been validated for TD muscle [Konno et al., 2021], so we set *s*_ECM_ = 1 for TD muscle (this corresponds to a value of 150 in Konno et al. [2021]). During our stress-strain tests, we then considered the possibility of the ECM component of muscle being stiffer (*s*_ECM_ = 1.33) and less stiff (*s*_ECM_ = 0.66). Note that a *s*_ECM_ value of 1.33 corresponds to a stiffness of 133 % compared to TD muscle, while a value 0.66 corresponds to a stiffness of 66 % relative to TD muscle. Data for the changes in stiffness of the ECM are not available experimentally, so these values of *s*_ECM_ were chosen to investigate the effects of altering this component. In summary, to investigate the effects of CP, our parameter ranges were *α* = 0.02 to 0.6, *c*_sarco_ = 0.0 to 0.75, and *s*_ECM_ = 0.66 to 1.33, while TD muscle had parameters *α* = 0.05, *c*_sarco_ = 0.0, and *s*_ECM_ = 1.0.

## 3 Results

### 3.1 Effects of *c*_sarco_

For TD muscle (*c*_sarco_ = 0.0, *s*_ECM_ = 1.0, and *α* = 0.05), we observed typical overall stress-stretch behavior for passive skeletal muscle: as the stress increased, the muscle stretch and sarcomere lengths also increased (Figure 2). The muscle block in its resting and stretched states is shown in Figure 2 C and D, respectively. The shift in the intrinsic passive sarcomere stress-stretch relationship, *c*_sarco_, affected the muscle behavior in both the stress-length for the sarcomeres and overall stress-stretch for the whole muscle relationships of the model (Figure 2). At *in vivo* lengths, we found that there is no longer zero stress for *c*_sarco_ > 0.0 (Figure 2 A). This indicates that larger forces are required to stretch muscles with increased sarcomere lengths, as well as to hold it at the resting length of the muscle. Additionally, optimal length of the sarcomeres no longer occured at the same resting length of the whole muscle. For the same range of stress values (0-3 × 10^5^ Pa), we saw a larger range of whole muscle stretches with larger *c*_sarco_ (Figure 2). This is likely due to effects from the base material and, therefore, the ECM, which acts to deform the muscle back to optimal length. At stretch values less than 1.0, the base material works to extend the muscle to optimal length, while the sarcomeres are still working to shorten the muscle for *c*_sarco_ > 0.0.

**Figure 2:**
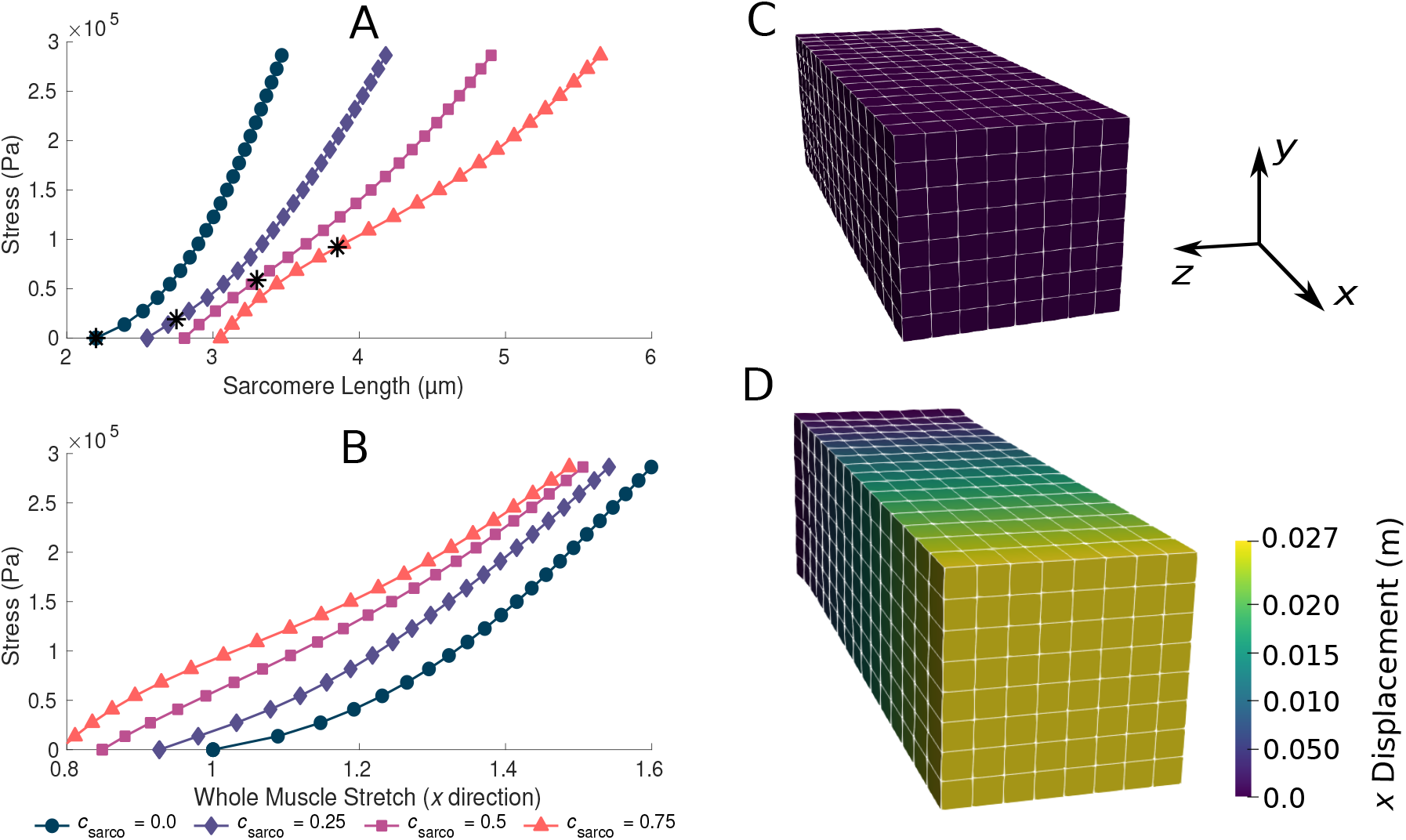
Plots of whole muscle stress in the along fibre direction against sarcomere length (A) and whole muscle stretch (B). Plots are from the computational model during passive lengthening with ECM volume fraction, *α*, of 0.05 and ECM stiffness factor, *s*_ECM_, of 1.0. Each line represents a shift in the sarcomere stretch by a factor of *c*_sarco_. * represents *in vivo* sarcomere length for corresponding *c*_sarco_. C and D show the mesh at resting length and at a deformed state after the stress has been applied to the model.

In addition to the behavior in the along-fibre direction of the muscle, *c*_sarco_ also affects the behavior transverse to the muscle fibres (Figure 3). We observed a similar change in concavity of the stress-stretch curves in both the stress-stretch relationships in the longitudinal (Figure 2 B) and transverse (Figure 3) directions. This demonstrates similar effects from the muscle ECM component in both directions. For stretch values in the *y* direction less than 0.85, the influence of the sarcomeres on the stress-stretch relationship decreased, and there was larger influence from the ECM. For smaller normal stresses in the longitudinal directions, there were larger effects from sarcomere length relative to larger stresses (Figure 3). The sarcomeres, acting only in the along-fibre direction, altered three dimensional deformation, which could effect muscle force production [Wakeling et al., 2020].

**Figure 3:**
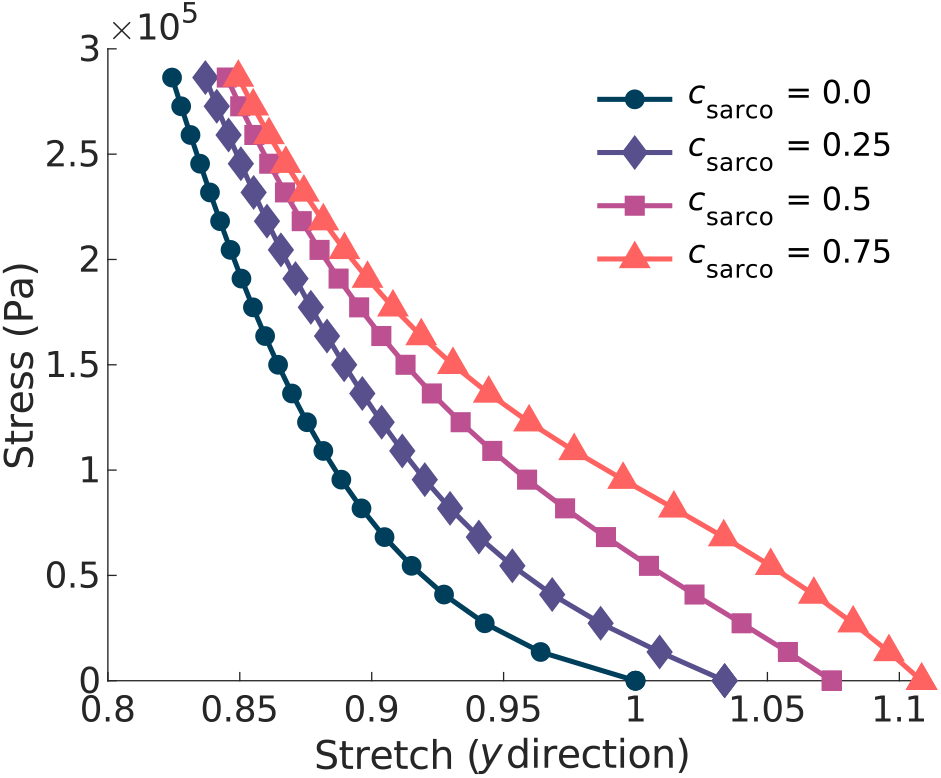
Plot of the normal stress applied in the along-fibre direction (*x*) against the stretch in the muscle transverse to the fibres. Given the symmetry in the muscle geometry transverse to the fibres (*y*), the stress-stretch response shown is the same in the *z* direction. Each line represents a shift in the sarcomere stretch by a factor of *c*_sarco_.

### 3.2 Effects of *α* and *s*_ECM_

The ECM properties also had a substantial effect on the overall stress-stretch relationships (Figure 4). Given the range of possible values for *α* (0.02-0.6), it had a larger effect on the muscle stiffness compared to *s*_ECM_. As *s*_ECM_ was increased, with *α* = 0.05 or 0.6, the overall stress-stretch relationship became more linear and covered a smaller range of stretch values. Increases in both *α* and *s*_ECM_ reduced the effect from the sarcomere length on the stress-stretch relationship (Figure 4). Overall, similar effects were observed from changing *s*_ECM_ and *α*, but given the change in composition of the base material, *α* had the greatest effect on the overall stress-stretch behavior.

**Figure 4:**
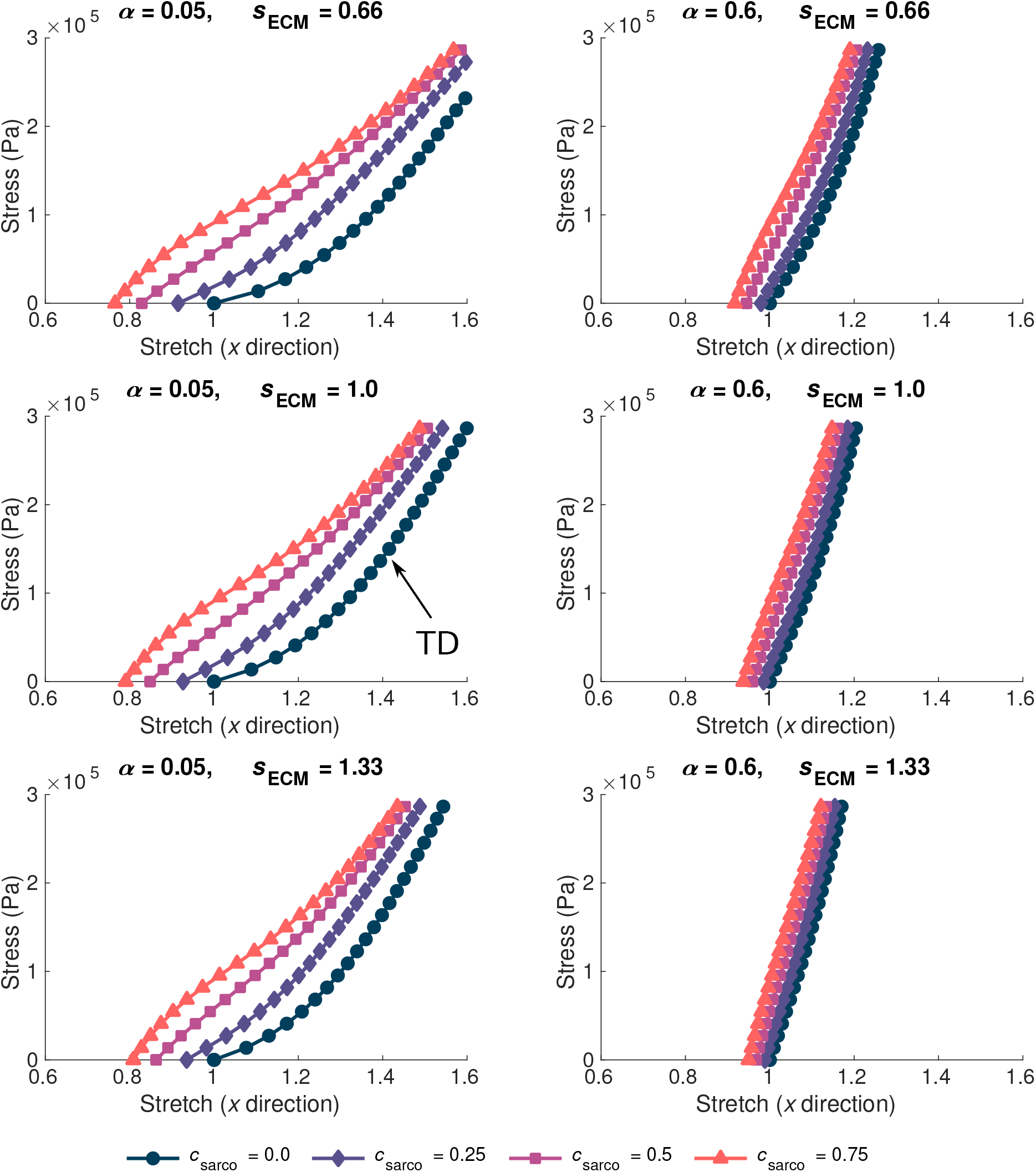
Stress-stretch plot during passive lengthening of the muscle model for various values of ECM volume fraction, *α*, and stiffness, *s*_ECM_. The traction was linearly increased on the *x* face of the muscle to 3 105 Pa, while the +*x* face was constrained in all directions. Individual lines on each plot represent a shift in sarcomere stretch by *c*_sarco_. Typically developed (TD) muscle has values of *α* = 0.05, *s*_ECM_ = 1.0, and *c*_sarco_ = 0.0, while cerebral palsy muscle could have a combination of *α* > 0.1, *s*_ECM_ = 0.66 or 1.33, and *c*_sarco_ > 0.

### 3.3 Muscle Stiffness

As *c*_sarco_ was increased up to a value of 0.5, the modulus of the muscle in the *x* direction increased at optimal length of the muscle (Figure 5 A,B). However, after a value of 0.5, the muscle modulus decreased, and this was observed when looking at the variations in *c*_sarco_ with constant *α* and *s*_ECM_ (Figure 5 A,B). The modulus at optimal length was dominated by *α*. For changes in *c*_sarco_ and *s*_ECM_, the change in modulus (at most 4 × 10^5^ Pa) was less than the possible variation in modulus with changes in *α* (up to 15 × 10^5^ Pa for highly fibrotic tissue). While holding *c*_sarco_ constant, we found a linear increase in the modulus when increasing *α* and *s*_ECM_ (Figure 5 C,D).

**Figure 5:**
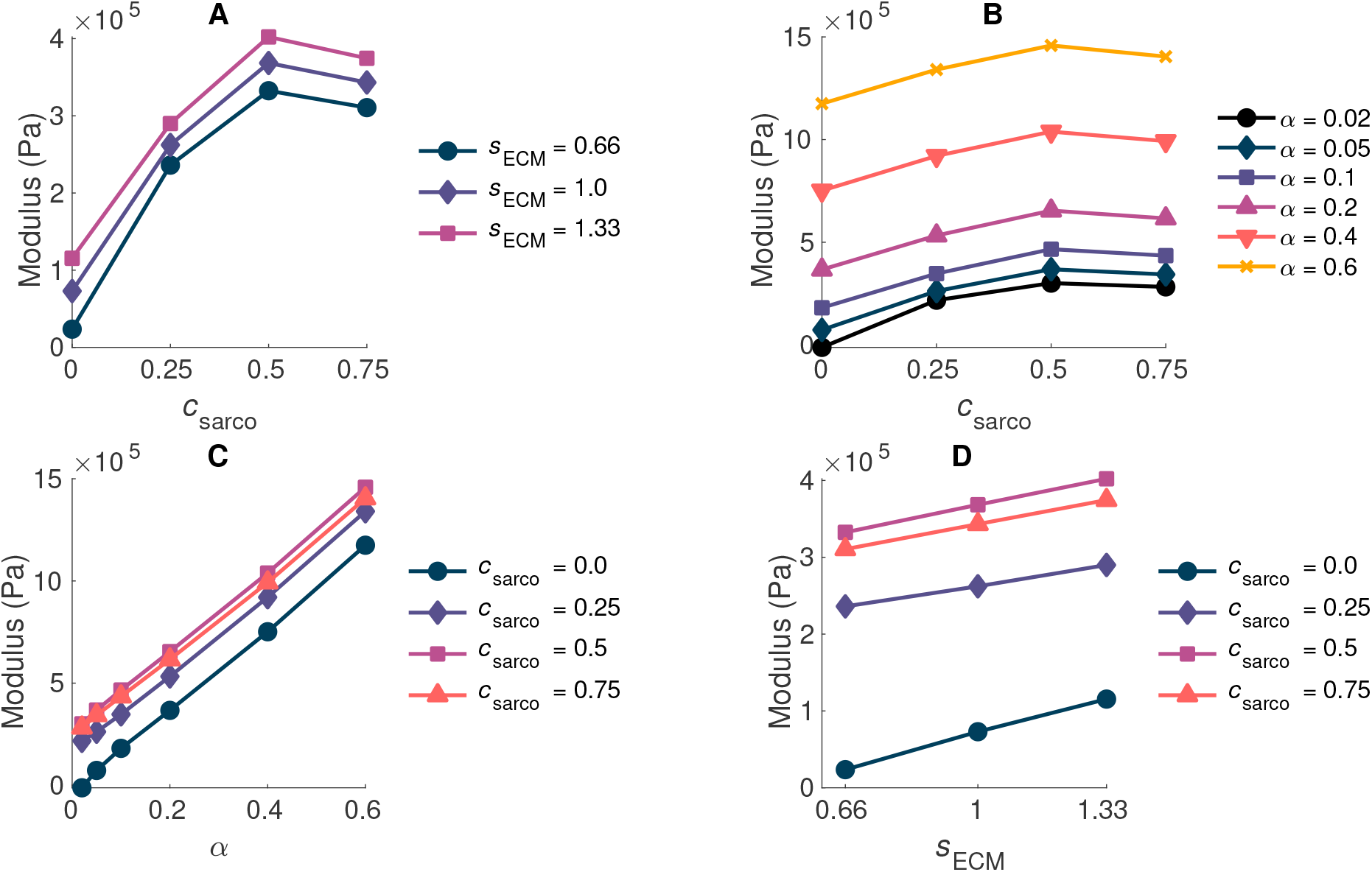
Plot of whole muscle modulus vs. *c*_sarco_ (A,B), *α* (C), and *s*_ECM_ (D) at optimal length 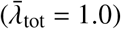.Where *α* is the ECM volume fraction, *s*_ECM_ is the ECM stiffness factor, and *c*_sarco_ is the shift in the sarcomere stretch. In plots A,D *α* is held constant at 0.05, and in plots B,C *s*_ECM_ is held constant at 1.0.

While holding *c*_sarco_ constant, there was a larger effect from volume fraction of the ECM, *α*, than the stiffness of the ECM, *s*_ECM_, on the overall muscle stiffness (Figure 6 A,B). However, as *α* was increased, there was a greater effect of *s*_ECM_. The nonlinear behavior, which showed increasing muscle modulus with increasing *α* and *s*_ECM_ was more pronounced at larger stretches (Figure 6 B,D). At a stretch of 1.20 in the *x* direction, the stiffness appeared to be more nonlinear when moving along the lines of constant *s*_ECM_ and when moving along the lines of constant *α* for *α* > 0.2 (Figure 6 C,D). When holding *s*_ECM_ constant, there was a larger effect *α* on the modulus of the muscle compared to *c*_sarco_, particularly at larger stretch values (Figure 6 C,D). As the stretch increased there was an increase in modulus from the ECM parameters; however, there was a decrease in the effects of *c*_sarco_ (Figure 6 E and F). The reduced influence of *c*_sarco_ was due to more pronounced behavior from the base material at larger stretches (Figures 4 and 6).

**Figure 6:**
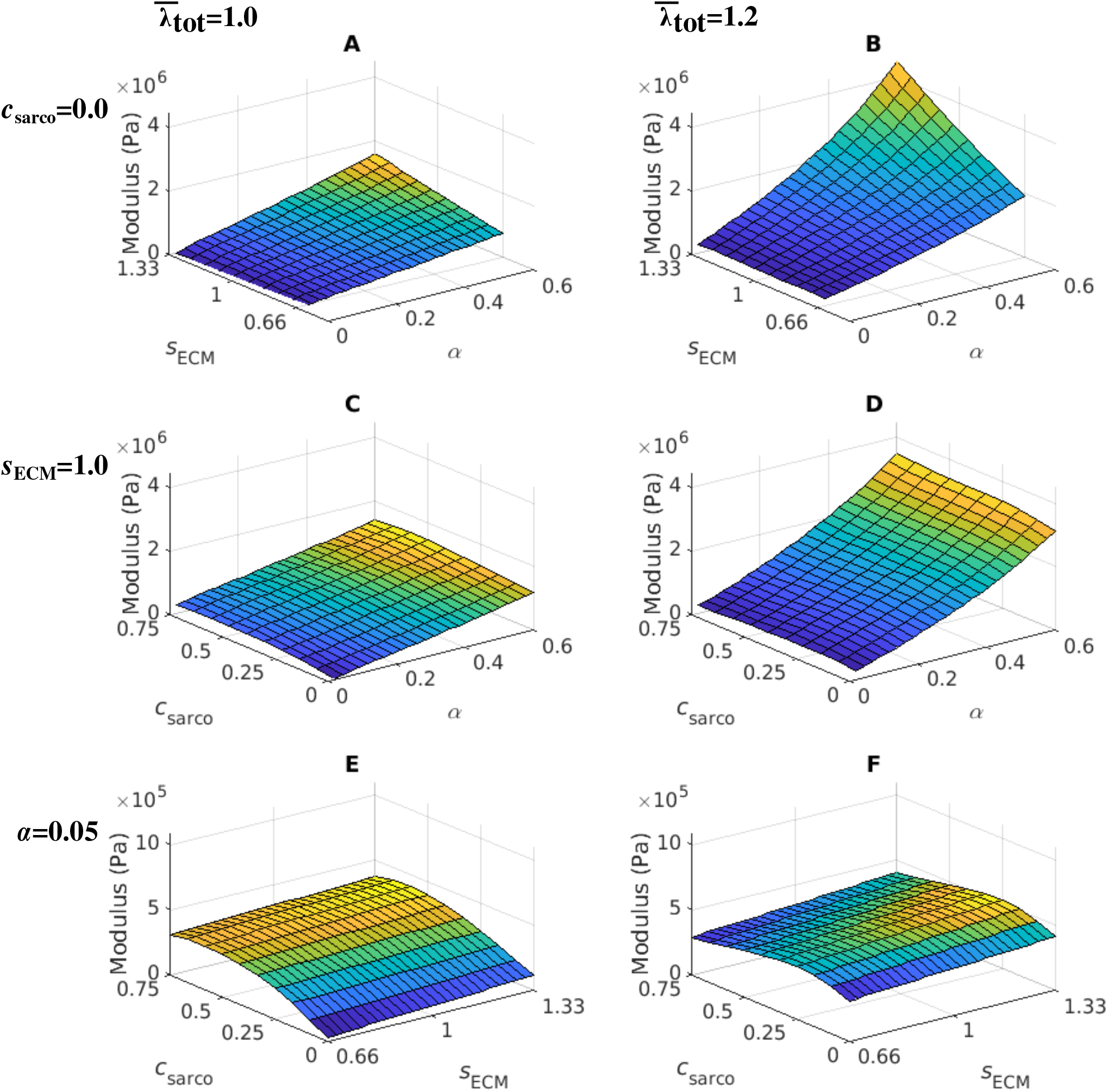
Surface plot of the whole muscle modulus at an average muscle stretch, 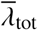, of 1.0 (A,C,E) and 1.2 (B,D,F). The ECM stiffness factor, *s*_ECM_, was varied between values of 0.66 and 1.33; ECM volume fraction, *α*, from 0.02 to 0.6; and shift in sarcomere stretch, *c*_sarco_, from 0.0 to 0.75. Modulus values were extracted from passive lengthening simulations with *c*_sarco_ = 0.0 in A,B; *s*_ECM_ = 1.0 in C,D; and *α* = 0.05 in E,F.

## 4 Discussion

In CP, alterations occur on the microstructural level that can influence whole muscle stiffness and reduce function. In particular, alterations to ECM properties and sarcomere length can occur; however, their relative contributions to muscle stiffness in CP is unknown. Isolating individual effects on passive muscle stiffness is difficult to do in experimental studies as there is large variability between subjects and individual muscles [Calvo et al., 2010, Mohammadkhah et al., 2016, Takaza et al., 2013]. Therefore, to determine whether the ECM properties or sarcomere lengths have more effect on the passive muscle behavior, we used a three dimensional continuum model [Konno et al., 2021, Rahemi et al., 2014, Ross et al., 2018, Wakeling et al., 2020]. This allowed us isolate the effects from individual microscopic components, and investigate the relative contributions to whole muscle function.

### 4.1 Physiological Changes to Muscle During Cerebral Palsy

The ECM is composed of a highly structured arrangement of collagen fibres and plays a substantial role in skeletal muscle mechanics [Gillies and Lieber, 2011]. In this study, we investigated the effects of changes to ECM volume fraction and stiffness on the whole muscle stiffness. Changes in ECM volume fraction have been observed in previous studies, particularly as a result of fibrosis (eg. Smith et al. [2011]). This corresponds to an increase in ECM material, while the contributions from the cellular components in muscle, such as the contractile fibres and other cells, decreases. In addition, fibrosis creates a physical barrier that can impact muscle regeneration [Chen and Li, 2009], which will reduce the ability for muscle to grow and add sarcomeres in the muscle fibres, further decreasing the compliance of the muscle. Additionally, it is possible that while the volume fraction stays constant, changes in the structure or composition of collagen types varies. However, studies have found that the ratio of collagen isoforms is the same in both TD and CP muscle [Smith et al., 2019], and so it is unlikely that a difference in collagen isoforms in muscle accounts for the increase in whole muscle stiffness with CP. It is possible that there are alterations in ECM structure, such as the organization of collagen fibres, that occur during CP [Lieber et al., 2003], and this could increase or decrease ECM stiffness depending on the change. Therefore, both effects were considered in the model to investigate the relative contributions to stiffness on a whole muscle level.

It has also been well documented that increases in the sarcomere length occur during CP [Lieber and Fridén, 2019, Tisha et al., 2019]. Stiffness changes have been reported on the fibre level by looking at the stress-strain relationship for TD and CP muscle fibres [Mathewson et al., 2014a]. It is possible that these effects are not only a result of increased sarcomere lengths, but due to different titin isoforms, as they could result in a different stress-strain relationship for the individual sarcomeres [Prado et al., 2005]. However, Smith et al. [2011] found that there is no change in the composition of titin isoforms between TD and CP muscle. Therefore, changes in stiffness due to the sarcomeres are likely an effect of increased stretches in the titin protein rather than different isoforms.

### 4.2 Microstructural Contributions to Whole Muscle Behavior

It has been demonstrated experimentally that the ECM has a significant contribution to muscle passive stiffness [Gillies and Lieber, 2011], and that fibrosis has been observed during CP [Lieber et al., 2003]. We found that the volume fraction of the ECM had a larger influence on whole muscle stiffness compared to ECM stiffness and sarcomere length. The contribution from the ECM increased as stretch increased (Figure 6), demonstrating a nonlinear relationship between the ECM volume fraction and muscle stretch. At larger stretch values, the ECM contributes more to the whole muscle stiffness; these nonlinear effects imply that fibrosis will substantially reduce the ability of a muscle to deform at larger stretch values. The ECM is composed of crimped collagen fibres, which likely do not contribute as much to the stress initially [Gillies and Lieber, 2011], and this is reflected in a smaller effect from the ECM volume fraction at optimal length. Currently, experimental data for the variation in stiffness of the ECM due to structural changes are not available; however, in the ranges tested in this study, the stiffness of the ECM did not alter whole muscle stiffness as much as the volume fraction.

The contribution of the sarcomere length to whole muscle stiffness varied depending on the ECM properties. There was minimal effect of the sarcomeres on the passive stiffness in the fibrotic tissue (Figure 5), which corresponds to volume fractions of ECM greater than 10 % and has be observed during CP [Smith et al., 2011]. Furthermore, the sarcomere effects are mitigated at larger stretches as the ECM begins to dominate the muscle stiffness. At whole muscle stretches near 1.0, we found a larger effect of sarcomere length (Figure 6), and this agrees with experimental data [Smith et al., 2011]. These findings demonstrate that if a muscle is operating near optimal length, then there might be a noticeable effect of sarcomere length. However, if the muscle has a larger range of motion, then the fibrotic effects would likely have a larger contribution to muscle passive stiffness. It is likely that the lengthening of the sarcomeres during CP has a larger effect on the active properties of the muscle (which we have not evaluated in this study) compared to the passive properties as noted by Lieber and Fridén [2019].

Using this model we can obtain a deeper understanding of the three-dimensional effects that occur in muscle altered by CP. As shown in previous modeling [Ryan et al., 2020, Wakeling et al., 2020] and experimental studies [Randhawa and Wakeling, 2018], the ability of a muscle to deform both in the along and transverse fibre directions can alter muscle function. In our model, the three-dimensional behavior is captured in part by the base material, which works to return the muscle to its original state. At longer muscle lengths, the base material will work in the same direction as the titin in the sarcomeres, which are trying to shorten the muscle. Meanwhile, for short muscle lengths (a muscle stretch less than one), the ECM will be working to expand the muscle back to optimal length. We have observed in the model that the stiffness of the muscle decreases after sarcomere lengths greater than 3.3 *μ*m (Figure 5), and this is due to the three dimensional behavior of the model we are using. In a one dimensional model, there are no effects from the volume conserving nature of the base material or other effects transverse to the fibres. This is a nearly incompressible and nonlinear model, and so the effects from the volumetric component of the model contribute more with larger shifts in the sarcomere force-length curve. While these effects have been observed based on our assumptions for the model (see Wakeling et al. [2020] and Konno et al. [2021]), which are typical of many finite element models [Blemker and Delp, 2005, Sharafi and Blemker, 2011, Spyrou et al., 2017], these effects have not been reported experimentally. Experimentally, the decrease in muscle stiffness may not be as substantial as the changes observed in this study; however it is likely a similar trend would appear. Another important consequence of the three-dimensional behavior is that changes occurring strictly in the along-fibre direction, such as changes in the sarcomere length, affect the stretch transverse to the fibres (Figure 3). In particular, the bulging and stretching in the transverse direction is decreased by increased *in vivo* sarcomere lengths, which increases the passive stiffness of the muscle fibres. Given this reduced movement in the transverse direction, it is likely that there would be a decreased contractile force produced given the significant effect of three dimensional deformation [Ryan et al., 2020]. This demonstrates that to accurately capture all of the effects from CP, investigating three dimensional behavior is required to completely understand the mechanical behavior of the muscle.

### 4.3 Model Parameters

Experimental studies are key to understanding the mechanical changes that occur during CP; however, many of the procedures are invasive and unable to determine the exact role each change due to CP plays in altering muscle stiffness [Lieber and Fridén, 2019, Smith et al., 2011, Tisha et al., 2019]. Additionally, there is contradicting data as to whether fibres, ECM, or both have a substantial contribution to passive stiffness [Mathewson et al., 2014a, Smith et al., 2011], which likely depends on the severity of the disease [Tisha et al., 2019]. There are less invasive procedures that have been developed to investigate the relationship between muscle stiffness and CP [Lee et al., 2016]; however, they are still unable to isolate the role of individual factors. For example, experimental studies have found that stiffness of CP muscle is twice as high as TD muscle [von Walden et al., 2017]; however, they were not able to determine which microstructural changes led to this increase in stiffness. While this model cannot directly determine which microstructural changes will cause this experimental increase in stiffness, it can provide insight into how various changes on the microscopic level could lead to these effects on muscle stiffness. We have observed that there is approximately double the increase in stiffness when the volume fraction of ECM in our model increases from 5 % to 20 %. Another possible way to achieve this increase in muscle stiffness is through increasing the stiffness of the ECM, or some combination of the two. The possible changes that cause increased stiffness can be investigated through our modeling approach and can indicate which factors may have the most impact on muscle behavior. It is difficult to perform experimental tests on whole muscles affected by CP, although tests have been done on mice with spasticity or fibrosis [Smith and Barton, 2014, Ziv et al., 1984], as muscle can only be dissected during surgery making it difficult to obtain data for a accurate comparison to similar TD muscle tissue. Muscle is a three-dimensional material, so applying a continuum model to CP muscle allows us to understand the underlying muscle mechanics. In particular, developing an understanding of the complete behavior of muscle will give insight into the role each micro-mechanical alteration that occurs during CP will play in whole muscle mechanical behavior.

While the model has the ability to investigate behavior of muscle that is difficult to examine experimentally, it relies on accurate experimental data for its intrinsic properties. Unfortunately, mechanical data for the effects of stiffness of the ECM are not available, so the value for the ECM stiffness parameter was chosen to vary by 33 % from healthy muscle. It is possible that changes in the structure of the ECM would change by more than this value; however, these values were chosen to probe the behavior of the ECM stiffness. Given the derivation of the whole muscle stress in the model (Equation 1) it is likely that the volume fraction of the ECM would still have the largest contribution to whole muscle stiffness. Both the ECM volume fraction and stiffness multiply the ECM stress response, so they have similar contributions to whole muscle behavior for small variations in their values. However, only the ECM volume fraction decreases the contributions from the cellular components. This aspect of the model is realistic, since it is not likely changes in the structure of the ECM will decrease the contribution of the fibres to whole muscle stiffness.

### 4.4 Limitations and Future Directions

A limitation of this model is the lack of current experimental certainty on changes that occur during CP. Many changes have been observed in CP; however, the extent to which structural changes occur are varied [Tisha et al., 2019]. Therefore, the effectiveness of the model in providing a comparison to CP muscle will depend on the specific muscle type. In the model, we have assumed for simplicity that with changes in the volume fraction of the ECM, there is no effect on the amount of force produced by the fibres. Any reduction in contribution from the muscle fibres is assumed to be included in the decrease in cellular component contribution to the base material response. We have not investigated the active behavior of muscle in this study, although it is fundamental in muscle function. In CP, the contractile force produced has been seen to decrease as the disease progresses [Stackhouse et al., 2005], so using this model to investigate the influence of ECM and sarcomere properties on active force would be valuable and would give additional insight into how the structural alterations that occur with CP individually impact muscle contraction. In this model, the properties of our TD muscle may not be representative of all muscles as the material properties vary both across and within studies [Calvo et al., 2010, Mohammadkhah et al., 2016, Takaza et al., 2013]. So, while the qualitative passive behavior is captured in this model, the exact values could vary between muscles. However, we would expect the general trends observed during this study to hold.

## 5 Conclusion

The purpose of this study was to determine the effects of the microstructural changes that are normally observed during experimental studies of CP muscle, including variation in ECM volume fraction, stiffness, and sarcomere length, on whole muscle stiffness. To do this, a three dimensional computational model of skeletal muscle was used, and overall stress-stretch relationships and muscle stiffness were calculated by measuring the passive stress of the muscle. We found that the volume fraction of the ECM had a larger effect on muscle stiffness compared to the ECM stiffness and sarcomere length, and that the effects of the sarcomere length were mitigated at larger ECM volume fractions. Investigating these effects provides deeper insight into diseased muscle mechanics, and will help to direct future experimental studies. In this study, we were able to determine the crucial role that the microstructural alterations observed in CP have on whole skeletal muscle behavior.

